# A Yeast-Based Screening System for Differential Identification of Poison Inhibitors and Catalytic Inhibitors of Human Topoisomerase I

**DOI:** 10.1101/2021.11.16.468530

**Authors:** Ahmed Seddek, Christian Madeira, Thirunavukkarasu Annamalai, Christopher Mederos, Purushottam B Tiwari, Aaron Z. Welch, Yuk-Ching Tse-Dinh

## Abstract

Inhibition of human topoisomerase I (TOP1) by camptothecin and topotecan has been shown to reduce excessive transcription of PAMP (Pathogen-Associated Molecular Pattern) - induced genes in prior studies, preventing death from sepsis in animal models of bacterial and SARS-CoV-2 infections. The TOP1 catalytic activity likely resolves the topological constraints on DNA that encodes these genes to facilitate the transcription induction that leads to excess inflammation. The increased accumulation of TOP1 covalent complex (TOP1cc) following DNA cleavage is the basis for the anticancer efficacy of the TOP1 poison inhibitors developed for anticancer treatment. The potential cytotoxicity and mutagenicity of TOP1 targeting cancer drugs pose serious concerns for employing them as therapies in sepsis prevention. The aim of this study is to develop a novel yeast-based screening system that employs yeast strains expressing wild-type or a dominant lethal mutant recombinant human TOP1. This yeast-based screening system can identify human TOP1 poison inhibitors for anticancer efficacy as well as catalytic inhibitors that can inhibit TOP1 DNA binding or cleavage activity in steps prior to the formation of the TOP1cc. In addition to distinguishing between such TOP1 catalytic inhibitors and TOP1 poison inhibitors, results from this yeast-based screening system will also allow elimination of compounds that are likely to be cytotoxic based on their effect on yeast cell growth that is independent of recombinant human TOP1 overexpression.

## 2. INTRODUCTION

Life-threatening complications and fatalities attributable to infections might not be directly caused by the infectious agent. Instead, they might happen as consequences of the host immune responses that are triggered by the infection process. Acute respiratory distress syndrome (ARDS) and pulmonary fibrosis (PF) are two deadly complications of coronavirus infection that are caused by the host immune response to the viral infection [1]. Bacterial infections can also lead to severe immune-related complications, such as ARDS, acute lung injury [2, 3], and TB sepsis which has an 80% mortality rate [4]. When microorganisms infect the host, pathogen-associated molecular patterns (PAMPs), which are distinct microbial structures and macromolecules, and damage-associated molecular patterns (DAMPs) that result from damaged host tissues, are identified by pattern recognition receptors (PRPs) that belong to the host immune system. The expression of PRPs through PAMP response genes activates the production of innate immune cells that target and kill the invading pathogens [5, 6]. However, excess immune response can cause systemic tissue injuries and sepsis [7]. Sepsis is defined as life-threatening organ dysfunction caused by a dysregulated host response [8] to infection caused by viruses, bacteria or other infectious agents. During sepsis, the overexpression of PAMP response genes leads to the uncontrollable release of inflammatory mediators, such as cytokines and chemokines leading to cytokine storm [9, 10].

Topoisomerases play a major role in resolving the topological problems encountered in transcription and replication [11-13]. Human TOP1 has been a target for clinically approved cancer treatment [14, 15]. FDA-approved TOP1 inhibitors are poison inhibitors derived from camptothecin [16, 17]. These poison inhibitors stabilize the enzyme-DNA covalent complex, preventing DNA religation and leading to cell death, which is useful in cancer treatment [15, 16, 18]. An earlier study showed that the activation of PAMP response genes requires human TOP1, and that the depletion and reversible inhibition of TOP1 selectively suppress PAMP response genes [19]. Treatment with camptothecin suppressed PAMP response genes, and rescued mice infected with *Staphylococcus aureus* from death by sepsis [19]. Topoisomerase activities can modulate chromatin structure and influence the gene expression patterns relevant for both cancer and inflammation [20]. A recent study also showed that therapy with the FDA-approved TOP1 poison inhibitor topotecan suppresses lethal inflammation induced by SARS-CoV-2 and protects hamsters against SARS-CoV-2-induced lethal inflammation [21]. However, cytotoxicity associated with topoisomerase poison inhibitors is not desirable in sepsis management. Catalytic inhibitors of human TOP1 that inhibits DNA binding and cleavage by TOP1 may be preferable because they are not expected to cause DNA breaks and cell death. Instead, such TOP1 catalytic inhibitors can potentially prevent DNA binding and cleavage to halt the catalytic cycle of TOP1, resulting in repressing PAMP response gene expression and alleviating the excess immune response caused by infections.

Currently, use of radio-labeled DNA substrate is generally needed to conduct the TOP1 DNA cleavage-religation inhibition assays to determine if a TOP1 inhibitor acts as a topoisomerase poison, or if the inhibitor acts as a catalytic inhibitor to prevent DNA cleavage [22, 23]. Yeast-based assays for identification of topoisomerase poison inhibitors are easier to perform and do not require handling of radioactive material. The budding yeast *Saccharomyces cerevisiae* has been used for several decades as a model to study human topoisomerases-targeting drugs [24, 25]. *S. cerevisiae* strains with null *top1* mutation and expressing recombinant human TOP1 were found to be sensitive to camptothecin [25], demonstrating that human TOP1 can be expressed and targeted successfully in yeast by poison inhibitors. A human TOP1 mutation, T718A, was reported to mimic the action of camptothecin, leading to the stabilization of the TOP1 covalent complex (TOP1cc), followed by accumulation of DNA breaks and cell death when the mutant TOP1-T718A enzyme is overexpressed under the control of the GAL1 promoter in yeast with *top1* null mutation [26, 27]. We propose that *S. cerevisiae* with *Δtop1* mutation that expresses galactose-inducible T718A mutant TOP1 can potentially be used as a screening system for catalytic inhibitors of human TOP1 that reduces the formation of the TOP1cc from the induced TOP1-T718A to rescue the growth of yeast cells overexpressing TOP1-T718A. Figure 1 illustrates this proposed yeast-based screening system. Recombinant wild-type human TOP1 forms only transient TOP1cc during relaxation reaction. A poison inhibitor like camptothecin will reduce the growth of *S. cerevisiae* cells when wild-type human TOP1 is overexpressed because of the accumulated DNA breaks. The mutant TOP1-T178A is lethal when overexpressed but cell growth can be rescued to some degree by catalytic TOP1 inhibitor that prevents the mutant TOP1-T718A from forming the covalent complex. This yeast-based screening system is validated in this study with a newly identified human TOP1 catalytic inhibitor.

**Figure 1.**
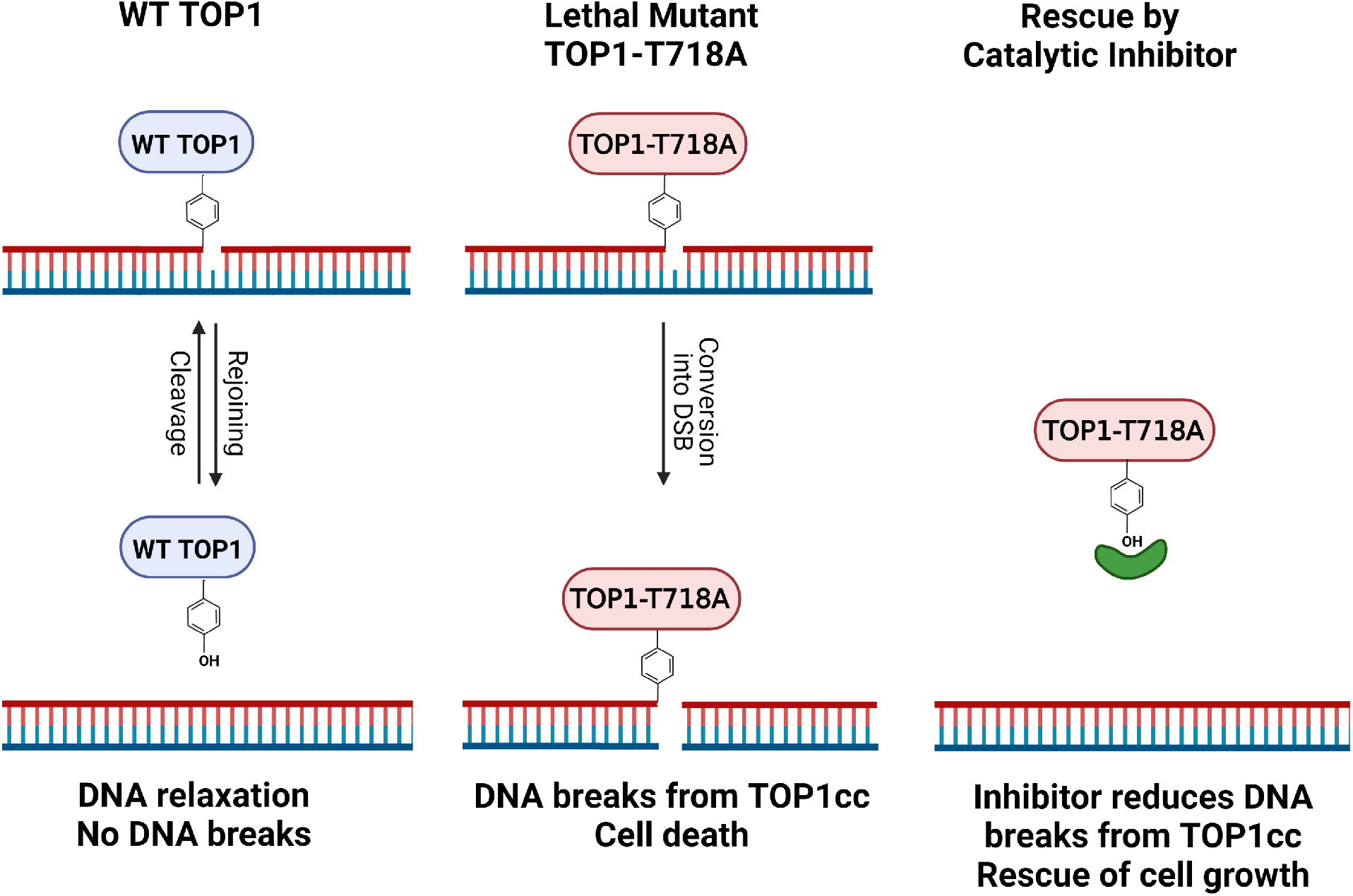
Scheme of yeast screening assay for human TOP1 catalytic inhibitors. Wild-type human TOP1 does not accumulate TOP1cc and DNA breaks. Increased DNA breaks from mTOP1cc because of T718A mutation leads to cell death. A catalytic inhibitor can reduce DNA breaks from mTOP1cc to rescue cell growth.

## 3. MATERIALS AND METHODS

### 3.1. Plasmid clones and yeast strains

The *S. cerevisiae* strain EKY3 (MATα, *ura3–52, his3*Δ*200, leu2*Δ*1, rp1*Δ*63, top1*::*TRP1*) previously described [28-30] was kindly provided by Dr. Leroy Liu at UMDNJ. The human TOP1 cDNA sequence (Accession No. NM_003286.3) with its starting ATG and termination codon was cloned into plasmid pYES2.1/TOPO (Invitrogen). The resulting clone pYES2-TOP1 expresses recombinant human TOP1 under the control of the GAL1 promoter. The T718A substitution was created in pYES2-TOP1 using the Q5 site-directed mutagenesis kit (New England BioLabs) with primers 5’-TCGCCTGGGAGCCTCCAAACTCA-3’ and 5’- ATCTGTTTATTTTCCTCTCGGTCTGTG-3’.

### 3.2. Enzyme and compounds

Recombinant human TOP1 enzyme was purchased from TopoGEN, and also purified from the lysate of *S. cerevisiae* EKY3/pYES2-TOP1 induced for expression of wild-type human TOP1with the PrepEase Histidine-Tagged Protein Purification Midi Kit – High Specificity (from Affymetrix) according to the manufacturer’s procedures. Camptothecin, myricetin and other flavonoids were purchased from Adooq Bioscience. NSC65860 was provided by NCI/DTP Open Chemical Repository.

### 3.3. Human TOP1 enzyme inhibition assay

Human TOP1 was assayed in reaction buffer containing 10 mM Tris-HCl pH 7.9, 1 mM EDTA, 0.15 M NaCl, 0.1% BSA, 0.1 mM spermidine, 5% glycerol [31]. The compounds were added to the enzyme suspended in the buffer, followed by addition of 200 ng of negative supercoiled DNA in the same buffer for a final volume of 20 μL. The amount of enzyme added was first determined to be the minimal amount required for complete relaxation of the input DNA following 30 min incubation at 37°C in the presence of 0.5 μl DMSO as control. The reactions were terminated with 4 μL of 6x SDS stop buffer (6% SDS, 6 mg/mL bromophenol blue, 40% glycerol) and was analyzed via electrophoresis in 1% agarose gel with TAE buffer [32]. The gel was stained in a 1 μg/mL solution of ethidium bromide for 30-45 minutes and rinsed for 10 -15 minutes in deionized water before photographed over UV light using the ProteinSimple AlphaImager Mini system. The percent inhibition was determined with the AlphaImager software based on intensity of supercoiled DNA remaining. The average and standard error of results from three experiments was plotted with GraphPad Prism to obtain the IC_50_ for inhibition.

### 3.4. Yeast growth assay

EKY3/pYES2-TOP1 and EKY3/pYES2-TOP1-T718A cultures were grown from plated colonies in Synthetic Complete – Uracil media (SC-U) supplemented with 1% glucose for 24 hours at 30°C. The culture was then diluted 1:50 into SC-U supplemented with 1% raffinose, and allowed to grow for a second day at 30°C. On the day of the experiment, OD_600_ values of the growing cultures were adjusted to 0.05 with SC-U supplemented with either 1% dextrose (expression repressor) or 1% galactose (expression inducer). The compounds tested were diluted serially in the same media in the successive rows of a sterile CELLTREAT polystyrene 96 well round bottom non-treated plate before the addition of an equal volume of the diluted cultures, achieving a final volume of 100 μL in each well. The plate was then incubated at 30°C for at least 66 hours. OD_600_ measurements were taken using a Synergy HT plate reader twice per day to monitor growth. Statistical significance of observed differences in OD_600_ from compound treatment was determined by using the two-way t-test for two independent means.

### 3.5. Preparation of yeast lysate for relaxation assay and western blot analysis

To confirm recombinant human TOP1 expression under the control of the GAL1 promoter in EKY3 cells, transformants were grown from colonies on plates in Synthetic Complete – Uracil (SC-U) media supplemented with 1% glucose for 24 hr at 30°C. The culture was then diluted 1:50 into SC-U supplemented with 1% raffinose, and allowed to grow for a second day at 30 °C. Finally, the OD_600_ values of the cultures were adjusted to 0.05 with SC-U media supplemented with either 1% dextrose (expression repressor) or 1% galactose (expression inducer) and allowed to grow for 66 hours at 30°C. Cell pellets were collected by centrifugation at 16,000 x g for 5 min.

For relaxation activity assay, cells were resuspended in a cell breaking buffer consisting of 50 mM sodium phosphate pH 7.4, 5% glycerol, 1 mM EDTA and 1 mM PMSF. OD_600_ values of the cell pellets in breaking buffer were adjusted to 25, then acid-washed glass beads were added to the buffer and vortexed with the resuspended cell pellets for four cycles of 30 sec agitation followed by 30 sec of cooling on ice. Soluble lysates were obtained by centrifugation at 16,000 x g for 10 min. Supercoiled plasmid DNA was used as substrate to assay the DNA-relaxing activity. Two microliters of soluble lysates were added to 200 ng of negatively supercoiled pBAD/Thio DNA in a reaction buffer that composed of 10 mM Tris HCl, pH 7.9, 1 mM EDTA, 150 mM NaCl, 0.1% BSA, 0.1 mM spermidine, and 5% glycerol [31]. All reactants were incubated for 30 minutes at 37°C, then each reaction was stopped by adding 6 μL of 4X SDS stop buffer (6% SDS, 0.3% bromophenol blue, 30% glycerol) and the supercoiled DNA and relaxed DNA were then separated by electrophoresis in 1% agarose gel. Purified recombinant human TOP1 (TopoGEN) was used as positive control.

For Western blot analysis of the human TOP1 expression, cell pellets were washed with 500 μL deionized water, then re-suspended in a volume of 0.1 M NaOH that resulted in OD_600_ readings of 2 – 3. Following 10 minutes at room temperature, the cell pellets were collected by centrifugation again and boiled for 5 minutes in the same volume of a sample buffer consisting of 60 mM Tris-HCl (pH 6.8), 4% 2-mercaptoethanol, 5% glycerol, 2% SDS, and 0.0025% bromophenol blue [33]. Samples were loaded and run in a 7.5% SDS-PAGE. After transferring separated proteins to a nitrocellulose membrane, the membrane was blocked with 5% Bovine Serum Albumin (BSA) at room temperature for 1 hour, then incubated with a 1:1000 (v/v) solution of mouse anti-human TOP1 antibodies (Developmental Studies Hybridoma Bank, Cat. No. CPTC-Top1-2) and mouse anti-Actin antibodies (Developmental Studies Hybridoma Bank, Cat. No. JLA-20) in 1X TBST buffer at 4 °C overnight. The membrane was washed three times for 5 min each in 1X TBST buffer, and then a 1:5000 (v/v) solution of horse-radish peroxidase (HRP) conjugated mouse IgG kappa binding protein (Santa Cruz Biotechnology Cat. No. sc-516102) in 1% BSA solution in 1X TBST buffer was incubated with the membrane at room temperature for one hour. After thorough washing and rinsing of the membrane in 1X TBST buffer, SuperSignal West Pico PLUS Chemiluminescent Substrate (Thermo Fisher Scientific) was used to visualize the protein bands.

### 3.6. Docking of NSC65860 to human TOP1

We performed docking of NSC65860 against a human TOP1 structure following procedures adopted in our previous publication [34]. Briefly, the 2D structure of NSC65860 (obtained from PubChem) was converted to a 3D structure and then a random 3D ligand conformer was obtained and converted to a pdbqt file by adding polar hydrogens using Open Babel [35]. PDB 1K4T [36] and I-TASSER [37] were used to generate the full length structure of human TOP1, followed by 10 ns molecular dynamics simulations using the NAMD simulation package [38]. The topology and parameter files used to generate and simulate the structure are as described previously [34]. Protein only coordinates from the covalent complex was selected for docking. AutoDockTools [39] was used to generate a pdbqt file of the human TOP1 structure. Finally, Autodock vina [40] was used to dock the NSC65860 against human TOP1 target structure. The dimensions of the search box was 90×120×130 Å^3^ that covered the entire human TOP1. Five independent dockings were performed using the same NSC65860 pdbqt file as mentioned above. The docked conformation with the highest affinity among the affinities of the top predicted model from the results of each docked result was then selected as the best conformation. VMD was used to generate the docked position of the best docked NSC65860 structure against human TOP1.

## 4. RESULTS

### 4.1 Inducible expression of human TOP1 in EKY3 yeast cells

To validate the catalytic function of recombinant human TOP1 induced in EKY3 yeast cells, EKY3 cell lysates were prepared and assayed with supercoiled plasmid DNA as substrate. The relaxation activity of the human TOP1 in the lysates were compared with purified recombinant human TOP1 (from TopoGEN) as control. As shown in Figure 2, EKY3/pYES2-TOP1 cells expressed active human TOP1 in the presence of galactose as inducer of the GAL1 promoter. The human TOP1 activity was present at only a very low level when the GAL1 promoter was repressed by dextrose. Soluble lysates of EKY3/pYES2-TOP1-T718A cells growth in presence of either dextrose or galactose did not show any relaxation activity as expected from previous characterization of this mutant TOP1 that is defective in DNA religation [26, 27].

**Figure 2.**
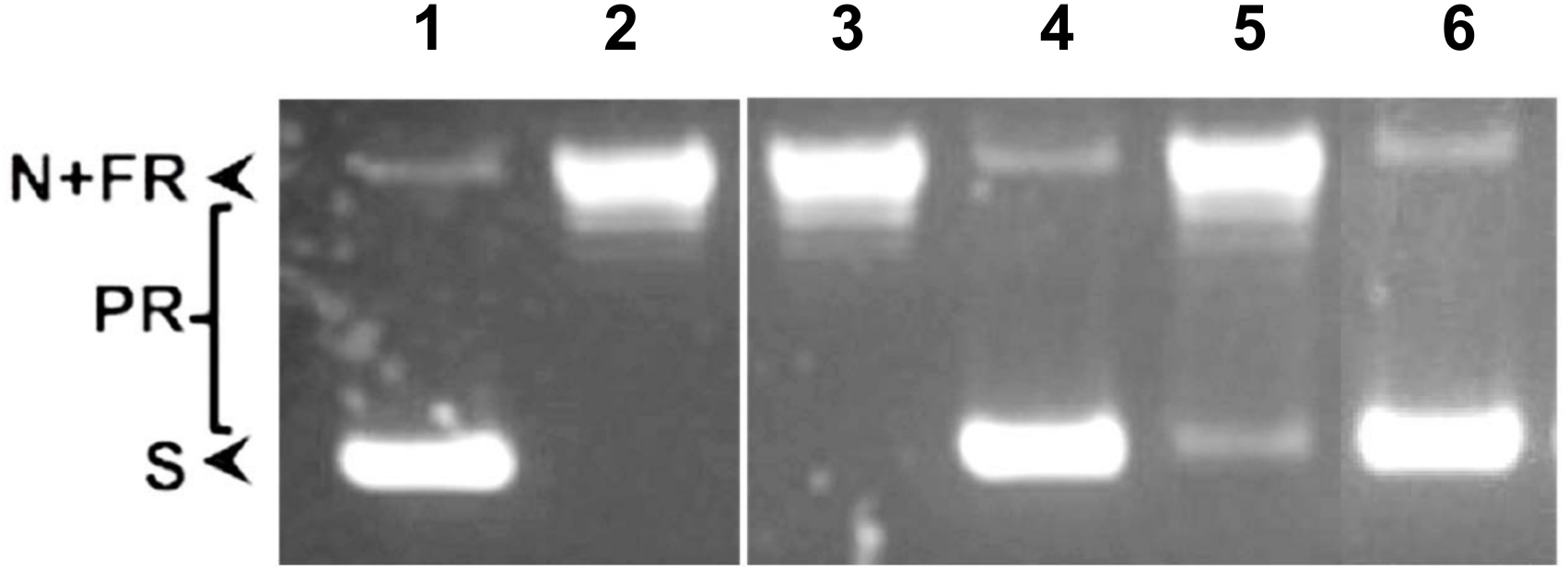
Activity of recombinant human TOP1 expressed in EKY3. Lane 1. Supercoiled plasmid DNA substrate. Lane 2. 1 U of human TOP1 from TopoGEN added. Lane 3 – 6. Equal amounts of lysates of EKY3 transformed with wild-type pYES2-TOP1 plasmid (lanes 3, 5) or mutant pYES2-TOP1-T718A plasmid (lanes 4, 6) were compared. Expression of recombinant human TOP1 was induced by galactose (lanes 3, 4) or repressed by dextrose (lanes 5, 6). S: Supercoiled DNA; N: nicked DNA; FR: Fully relaxed DNA; PR: partially relaxed DNA. The lanes shown here are from the same gel.

### 4.2. Growth inhibition of EKY3 yeast cells from induction of mutant TOP1-T718A

We confirmed that the accumulation of the covalent intermediate formed by recombinant TOP1-T718A in EKY3 led to galactose-dependent growth inhibition. Growth observed from serial dilutions spotted on plates (Figure 3A) showed equal growth of wild-type and mutant transformants on plate with dextrose but little or no growth for transformant of mutant TOP1-T718A on plate with galactose. The OD_600_ of cultures (Figure 3B) also showed growth inhibition of EKY3 from galactose-induced expression of mutant TOP1-T718A topoisomerase while galactose induction of wild-type TOP1 is well tolerated.

**Figure 3.**
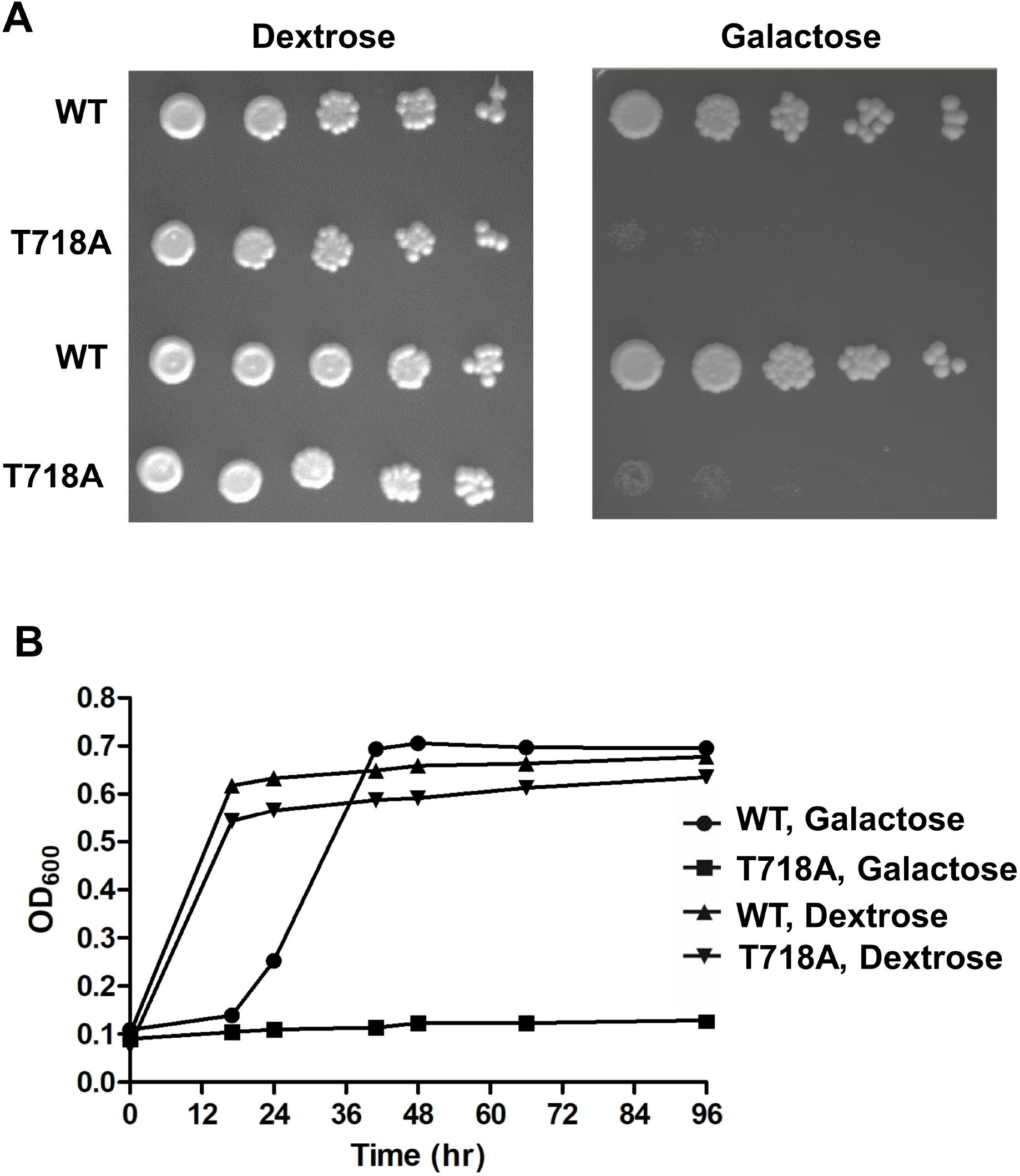
Galactose-dependent growth inhibition of EKY3 from expression of TOP1-T718A mutant. (A) EKY3/pYES2-TOP1 and EKY3/pYES2-TOP1-T718A saturated cultures in SC-U media with 2% raffinose were diluted to OD600 = 0.5 before 5-fold serial dilutions were spotted on SC-U agar plates supplemented with either 2% dextrose or galactose. (B) Growth curves of EKY3/pYES2-TOP1 and EKY3/pYES2-TOP1-T718A in SC-U media supplemented with 1% dextrose or galactose.

### 4.3. Effect of camptothecin on the growth of EKY3/pYES2-TOP1 and EKY3/pYES2-TOP1-T718A

Camptothecin is an extensively characterized poison inhibitor of human TOP1 with synthetic analogs that are used clinically in anticancer treatment [14, 17]. Addition of camptothecin to culture of EKY3/pYES2-TOP1 resulted in growth inhibition to a much greater extent in the presence of galactose versus dextrose, as expected from the toxic effect from the trapping of the covalent complex formed by induced recombinant TOP1 (Figure 4). In contrast, camptothecin had relatively little effect on growth of EKY3/pYES2-TOP1-T718A with expression of the mutant TOP1-T718A induced by galactose, since this mutant protein already forms a stabilized TOP1cc. These results demonstrate that utility of the yeast screening system for studying the TOP1 poison inhibitors.

**Figure 4.**
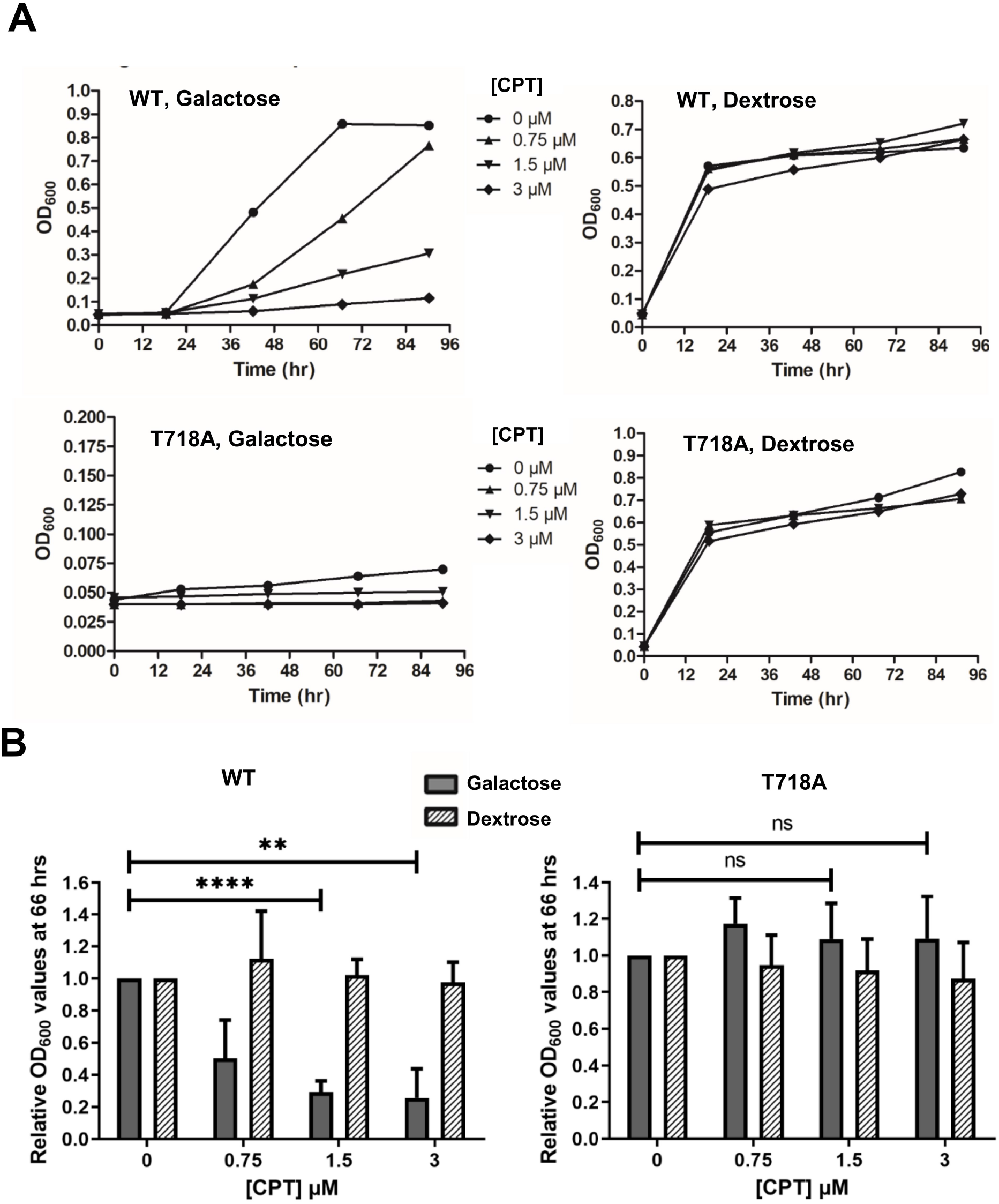
Effect of camptothecin on growth of EKY3/pYES2-TOP1 and EKY3/pYES-TOP1-T718A in media with dextrose or galactose. (A) Representative growth curves with increasing concentrations of camptothecin present. (B) Effect of camptothecin on relative OD600 of cultures after 66 hrs in dextrose media with non-induced versus galactose media with induced expression of wild-type (WT) or T178A mutant recombinant human TOP1. ****: P < 0.0001, **: P < 0.01, ns: not significant.

### 4.4. Characterization of natural product TOP1 inhibitor with yeast-based assay

In addition to camptothecin, natural products have been reported in the literature as human TOP1 inhibitors [41-44]. It is not always clear if these natural products act as poison inhibitors or catalytic inhibitors in their TOP1 inhibition mechanism [44]. A set of nine flavonoids and similar compounds (Figure S1) were first tested for inhibition of TOP1 relaxation activity at up to 250 μM concentration. Myricetin was the only compound found in this preliminary study to inhibit the relaxation activity of recombinant human TOP1 expressed in yeast completely. Follow-up assay of serial dilution of myricetin showed that the IC_50_ for inhibition of more highly purified human TOP1 from TopoGEN is ∼10 μM (Figure 5A). Testing of myricetin in the yeast-based assay (Figure 5B) showed that it inhibited growth of EKY3/pYES2-TOP1 with either galactose or dextrose in the media, with a greater degree of growth inhibition in galactose containing medium. There is also no increase in the growth of EKY3/pYES2-TOP1-T718A. These results suggested that in addition to inhibiting TOP1 activity as a poison inhibitor, myricetin can inhibit yeast cell growth with TOP1-independent mechanism. Myricetin was found previously to inhibit both human TOP1 and TOP2α [41, 43]. The yeast-based screening assay described here is thus useful for identifying TOP1 inhibitors that may have other cellular targets.

**Figure 5.**
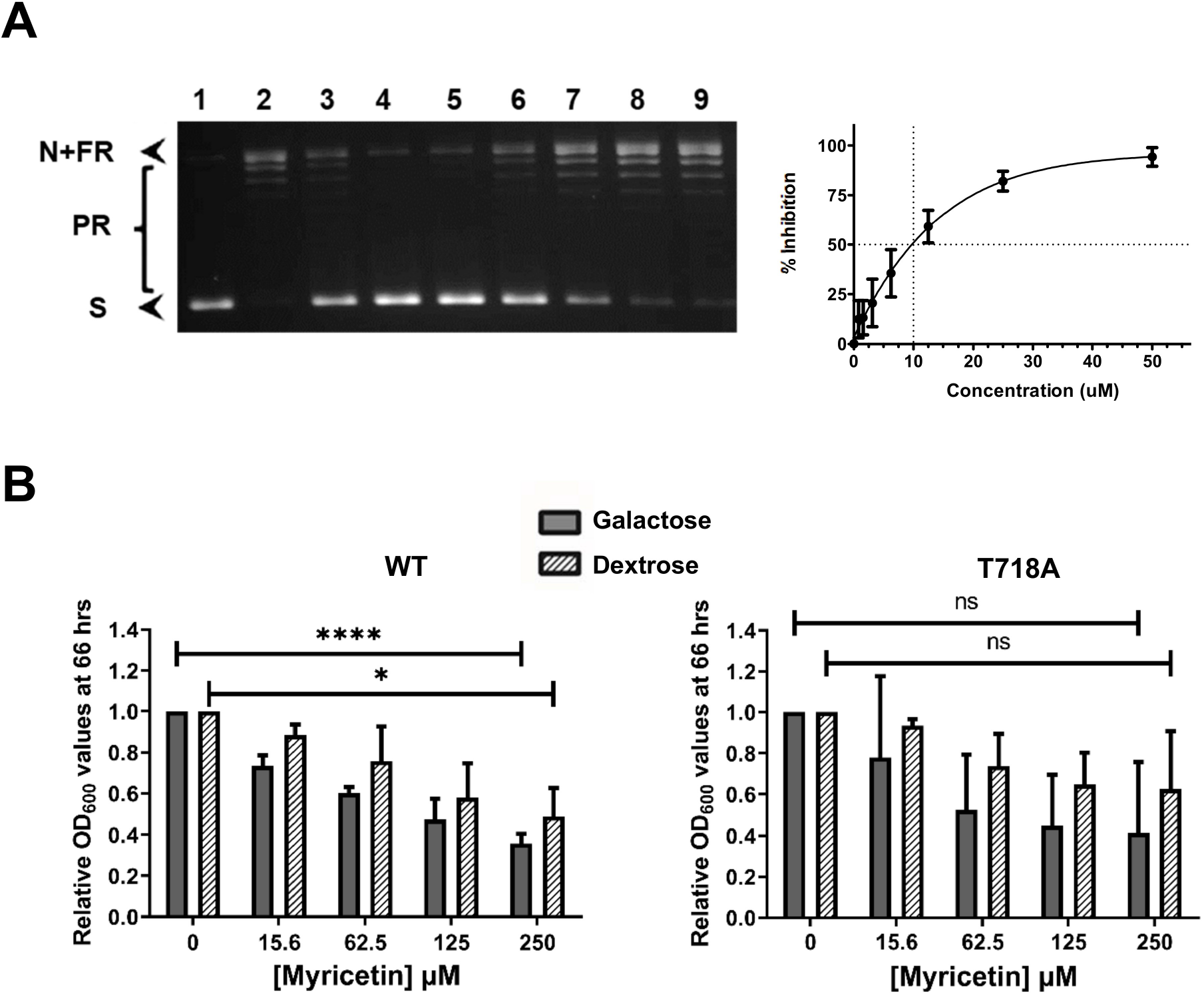
Testing of myricetin in human TOP1 relaxation and yeast-based growth assay. (A) Human TOP1 (0.3 U from TopoGEN) was assayed for relaxation of supercoiled plasmid DNA in the presence of serial dilutions of myricetin. Lane 1: no enzyme; lane 2: DMSO control; lane 3: 25 μM CPT; lanes 4 – 9: 50, 25, 12.5, 6.2, 3.1, 1.6 μM myricetin. S: Supercoiled DNA; N: nicked DNA; FR: Fully relaxed DNA; PR: partially relaxed DNA. Results from three replicates were plotted to obtain the IC_50_. (B) Effect of myricetin on relative OD_600_ of cultures after 66 hrs in 1% dextrose media with non-induced or 1% galactose media with induced expression of wild-type (WT) or T178A mutant recombinant human TOP1. ****: P < 0.0001, *: P < 0.05, ns: not significant.

### 4.5. Characterization of a new TOP1 catalytic inhibitor candidate

Compounds with functional groups that may compete for nucleic acid binding to topoisomerases were obtained from NCI Developmental Therapeutics and tested for human TOP1 inhibition in relaxation assays as well as the yeast-based screening assay described here. NSC65860 was found to be a strong inhibitor of human TOP1 relaxation activity, with IC_50_ of 0.89 μM for inhibition of the relaxation activity (Figure 6A). NSC65860 had no significant effect on the growth of EKY3/pYES2-TOP1 in the presence of either dextrose or galactose (Figure 6B). In the presence of dextrose to repress the expression of the toxic mutant TOP1-T718A protein, NSC65860 had also little effect on growth of EKY3/pYES2-TOP1-T718A However the growth of EKY3/pYES2-TOP1-T718A in the presence of galactose was enhanced significantly by the addition of NSC65860, with up to 3.5-fold increase in OD_600_ at 66 hr. (Figure 6B). This is consistent with NSC65860 acting as TOP1 catalytic inhibitor to prevent the formation of intracellular TOP1-T718A covalent complex from the induced toxic mutant TOP1.

**Figure 6.**
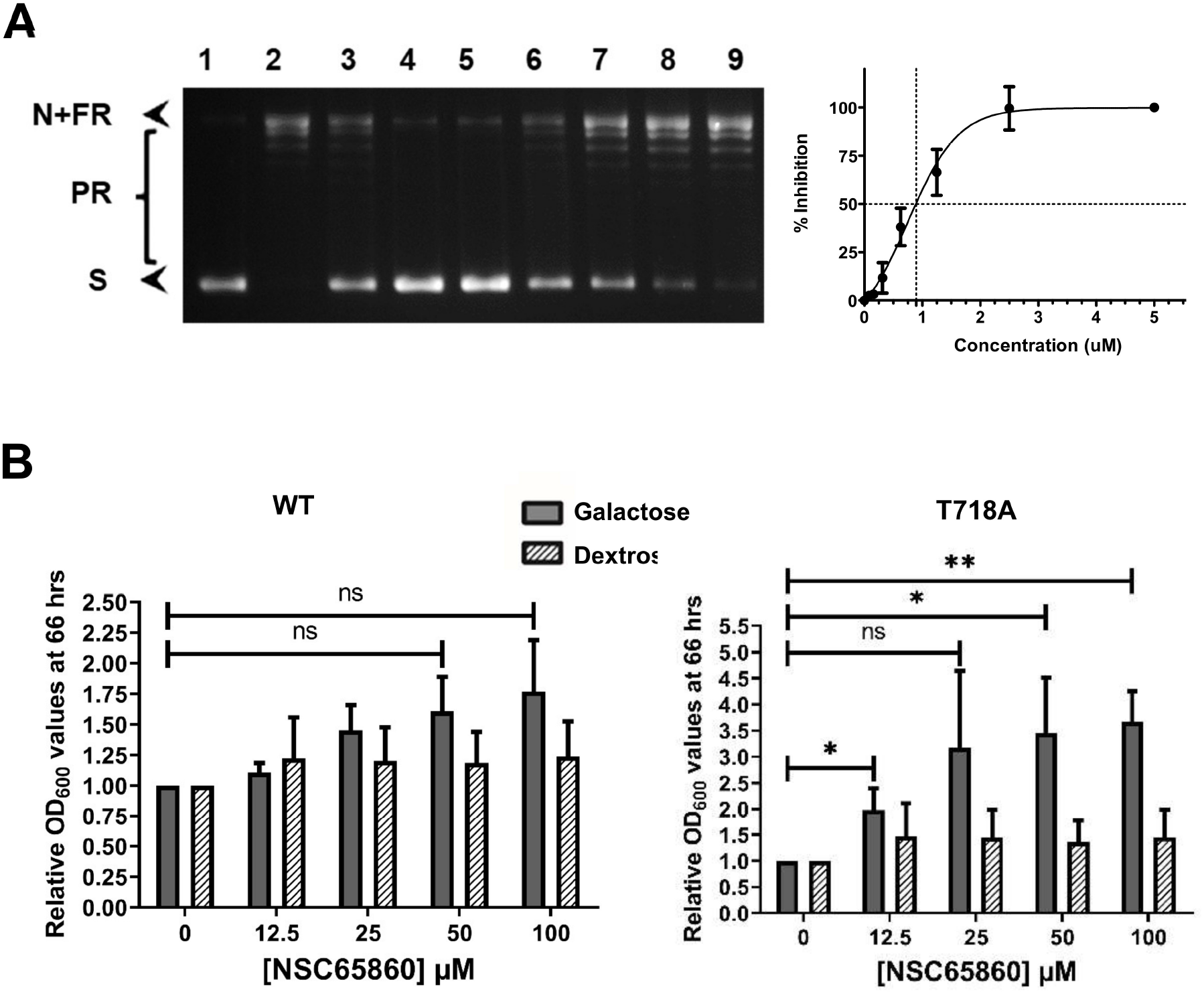
Testing of NSC65860 in human TOP1 relaxation and yeast-based growth assay. (A) Human TOP1 (0.3 U from TopoGEN) was assayed for relaxation of supercoiled plasmid DNA in the presence of serial dilutions of NSC65860. Lane 1: no enzyme; lane 2: DMSO control; lane 3: 25 μM CPT; lanes 4 – 9: 5, 2.5, 1.2, 0.62, 0.31, 0.16 μM NSC65860. S: Supercoiled DNA; N: nicked DNA; FR: Fully relaxed DNA; PR: partially relaxed DNA. Results from three replicates were plotted to obtain the IC_50_. (B) Effect of NSC65860 on relative OD_600_ of cultures after 66 hrs in 1% dextrose media with non-induced or 1% galactose media with induced expression of wild-type (WT) or T178A mutant recombinant human TOP1. * P < 0.05; ** P < 0.01; *** P < 0.001

It needs to be confirmed that treatment of EKY3 with NSC65860 did not impair the induction of recombinant protein expression from the GAL1 promoter. Western blot with antibodies against human TOP1 was used to show that treatment with as much as 100 μM NSC65850 had little or no effect on the production of recombinant human TOP1 from the GAL1 promoter following induction with 1% galactose (Figure 7). Even though induction of TOP1-718A cannot be examined directly due to the lethal effect of the mutant protein, the results for induction of WT TOP1 support the proposed role of NSC65860 in preventing the formation of lethal TOP1 covalent complex as mechanism of enhancing growth of EKY3/pYES2-TOP1-T718A in galactose medium.

**Figure 7.**
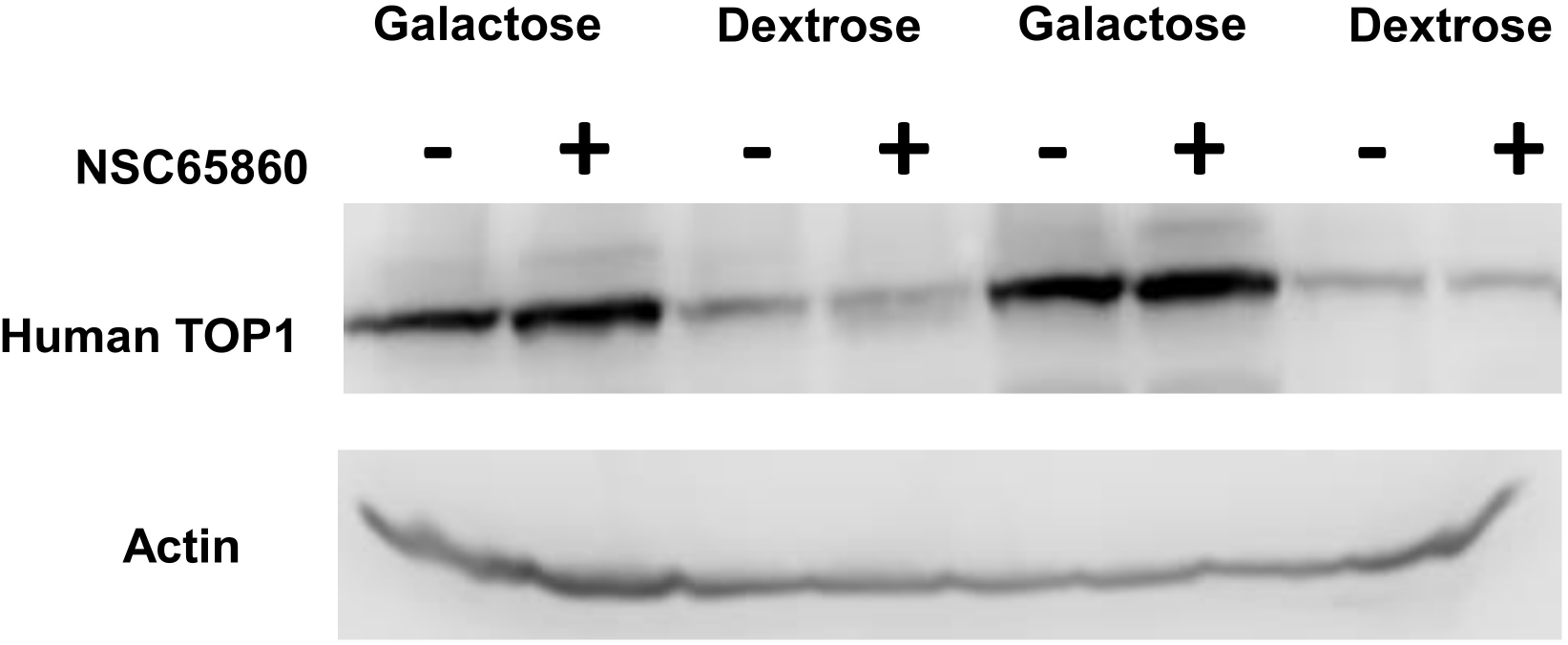
Western blot analysis of effect of NSC65860 on induction of GAL1 promoter. Total lysates of equal number of EKY3/pYES2-TOP1 cells grown in media with 1% galactose or dextrose, with and without treatment of 100 μM NSC65860 were analyzed in duplicates by SDS-PAGE followed by western blots with antibodies against human TOP1 and actin.

### 4.6. Docking of NSC65860 near human TOP1 active site

Molecular docking is a widely used computational technique to predict ligand-receptor complexes. In this work, we used Autodock vina to predict a possible binding site for NSC65860 when human TOPO1 was used as a target. The structure of NSC65860 is shown in Figure 8A. Notably, the search box used in our docking procedure covered the entire TOP1 structure allowing NSC65860 to dock randomly in the target. The human TOP1 target structure is taken from its covalent complex with DNA. Figure 8B shows the best-predicted docked position of NSC65860 in human TOP1. Our result predicts that the region near the active site tyrosine (Y732) is a possible binding site for NSC65860.

**Figure 8.**
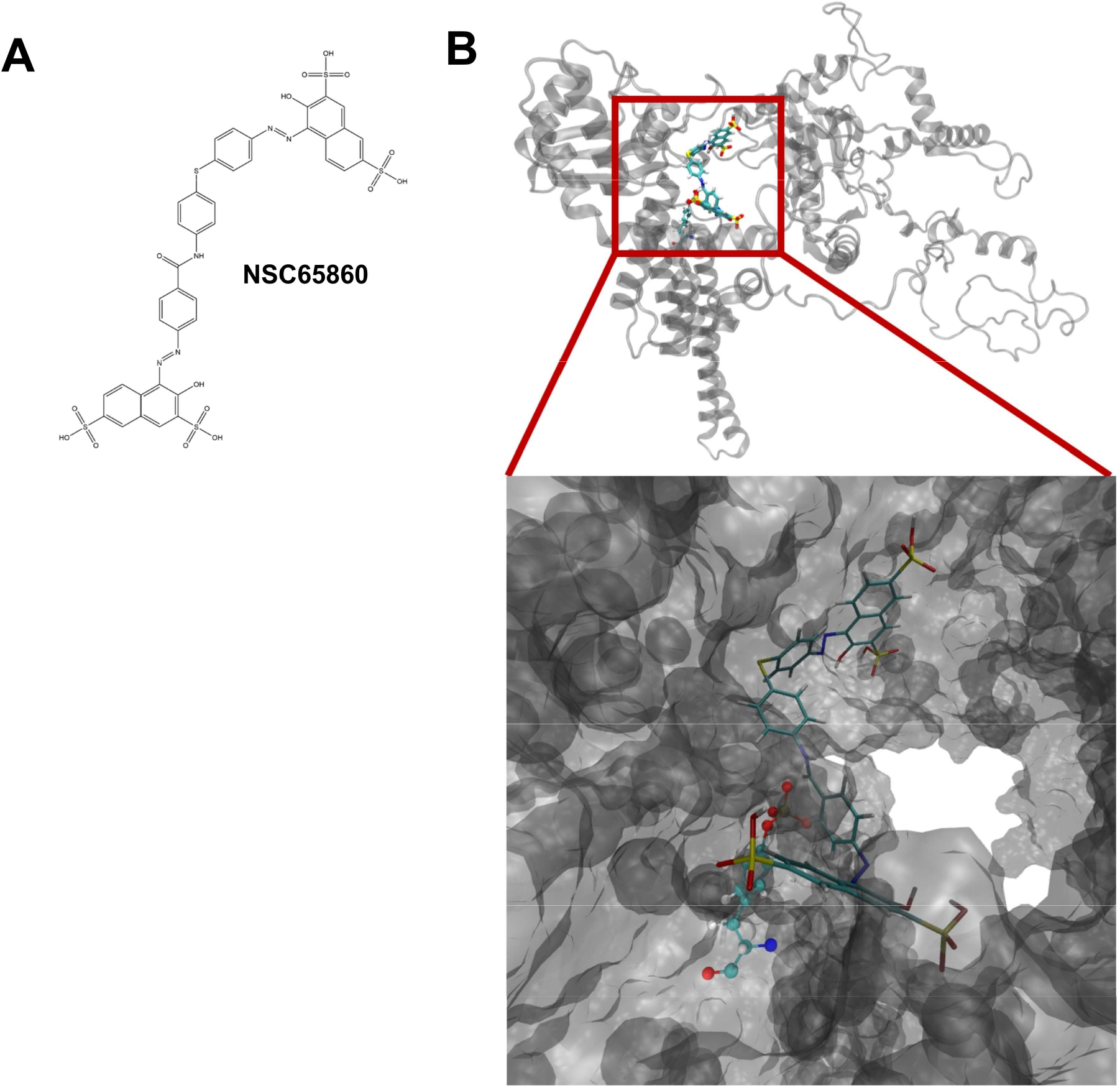
Docking of NSC65860 near the active site of human TOP1. (A) Structure of NSC65860. (B) Docked position of NSC65860 against human TOP1 target. The structure in grey color represents human TOP1. CPK structure represents Y723 in the active site, and licorice structure represents NSC65860. Upper panel shows the entire human TOP1 with the docked NSC65860. Lower panel shows the TOP1-NSC65860 complex zoomed near the active site Y723.

## 5. DISCUSSIONS

The lethal mutant human TOP1-T718A enzyme forms a stabilized covalent complex following DNA cleavage, leading to accumulation of DNA breaks and eventually cellular death when overexpressed from the GAL1 promoter in yeast cells cultured in media containing galactose [27]. *S. cerevisiae* cells overexpressing TOP1-T718A can be used as a model to screen for catalytic inhibitors of human TOP1, because a catalytic inhibitor can prevent DNA binding or cleavage by human TOP1-T718A to reduce the accumulation of TOP1cc with cleaved DNA, which would increase the viability of these cells when grown in media with galactose inducer. The TOP1 catalytic activity is not required for yeast cell growth. A specific human TOP1 inhibitor should not negatively impact the growth rates of yeast cells expressing WT human TOP1. In this study, we used yeast EKY3 transformed with WT or T718A mutant TOP1 clones to demonstrate that compound NSC65860 behaves as expected for a catalytic inhibitor of human TOP1 in this yeast-based assay. The growth of EKY3/pYES-TOP1-T718A in the presence of galactose was enhanced by >3.5-fold while growth in the presence of dextrose was not changed. NSC65860 also did not affect growth of EKY3/pYES-TOP1 in either galactose or dextrose containing media. Western blot analysis of soluble lysates of NSC65860-treated EKY3 cells showed that it does not affect the expression of recombinant human TOP1 from the GAL1 promoter, in support of its action as a catalytic inhibitor that inhibits DNA binding or cleavage by human TOP1. Additional research is required to determine the exact mechanism of action of NSC65860 and its impacts on PAMP gene expression in human cells, so that it can potentially be optimized into a candidate for sepsis treatment.

The results reported here provide validation that this yeast-based screening system could be used as a rapid and economical screening approach for identifying specific catalytic inhibitors of human TOP1 that are non-cytotoxic. Compared to enzyme-based gel electrophoresis assay, the yeast-based assay would require less resources to scale up for screening of a large library of molecules for TOP1 inhibitors. Candidates for either TOP1 catalytic or poison inhibitors can be identified based on the differential effects of the compound on growth of EKY3/pYES-TOP1 and EKY3/pYES-TOP1-718A in media containing dextrose versus galactose. A specific human TOP1 poison inhibitor would diminish growth only when WT TOP1 expression is induced by galactose. Myricetin resulted in most significant growth suppression of EKY3 with galactose induction of WT TOP1 to indicate that it can act as a TOP1 poison inhibitor. There was also some degree of suppression of growth of EKY3/pYES-TOP1 in the presence of dextrose, suggesting that myricetin can affect yeast cell viability in mode of action that does not involve TOP1 inhibition. Significant growth suppression EKY3/pYES-TOP1 in dextrose containing media can thus be an indication that the molecule is not likely to be a selective human TOP1 inhibitor and may not be a good candidate for consideration for further investigation of TOP1 poison or catalytic inhibitors for potential clinical applications. This useful information on selectivity at the cellular level can also be obtained readily from this yeast-based screening system.

## Acknowledgment

We thank Norhan Mohammed for technical assistance.

## Abbreviations

PAMP: pathogen-associated molecular pattern
ARDS: acute respiratory distress syndrome
PRP: pattern recognition receptor
TOP1: Topoisomerase I
TOP1cc: TOP1 covalent complex
CPT: camptothecin
WT: wild-type

---

**Figure S1.**
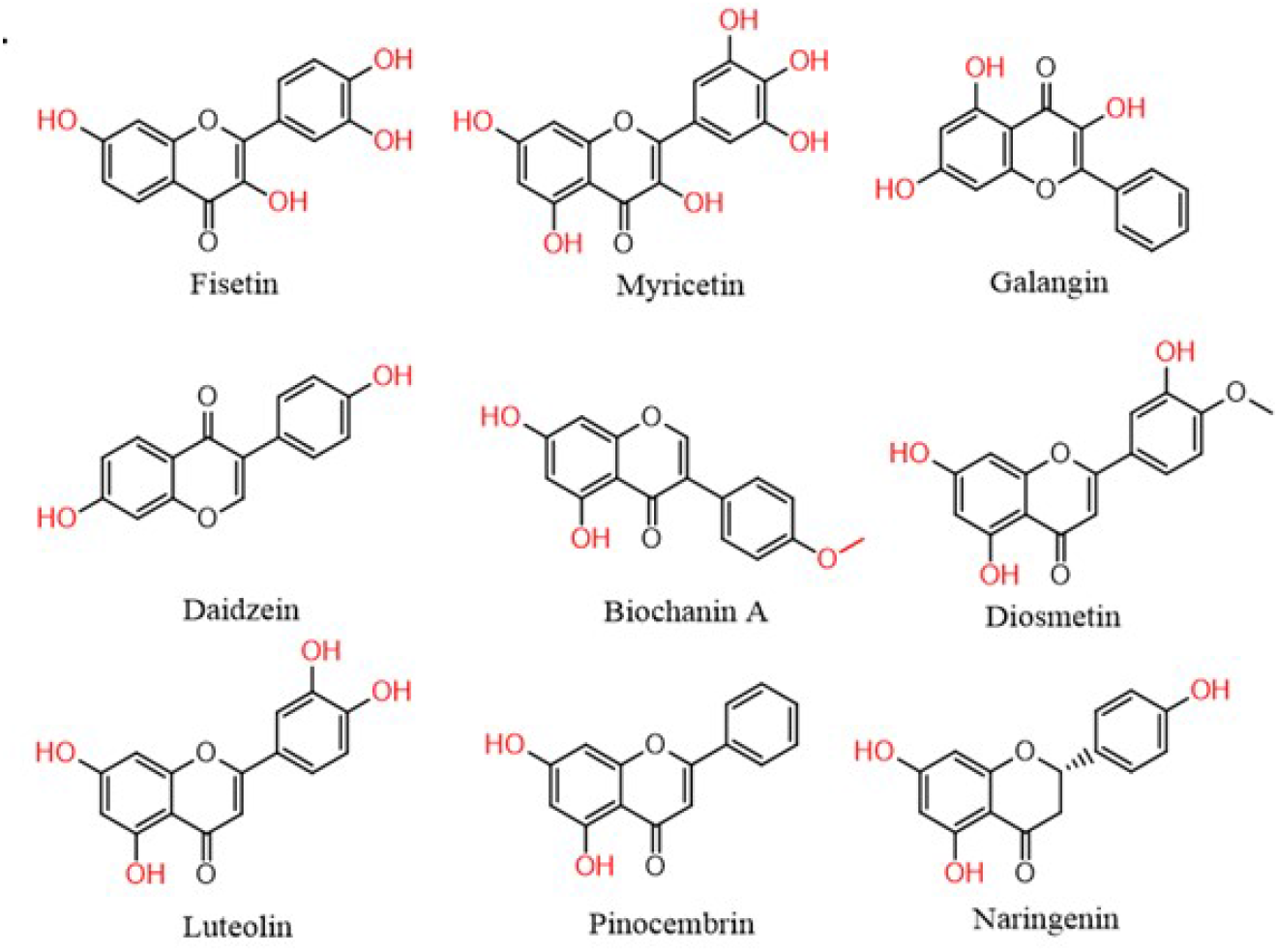
Natural products tested for inhibition of TOP1 relaxation activity.

## Notes

**Conflict of interest** The authors declare no conflict of interest

### Competing Interest Statement

The authors have declared no competing interest.

